# Voluntary actions modulate perception and neural representation of action-consequences in a hand-dependent manner

**DOI:** 10.1101/2020.01.12.903054

**Authors:** Buaron B., Reznik D., Gilron R., Mukamel R.

**Affiliations:** Sagol School of Neuroscience and School of Psychological Sciences, Tel-Aviv University, Tel Aviv, Israel; Department of Psychology, Center for Brain Science, Harvard University, Cambridge, MA, USA; Department of neurological surgery, UCSF School of Medicine, UCSF, San Francisco CA

## Abstract

Evoked neural activity in sensory regions, and perception of sensory stimuli, are modulated when the stimuli are the consequence of voluntary movement as opposed to an external source. It has been suggested that such modulations are due to efference copies of the motor command that are sent to relevant sensory regions during voluntary movement. Given the anatomical-functional laterality bias of the motor system, it is plausible that the pattern of such behavioral and neural sensory modulations will exhibit a similar bias, depending on the effector that was used to trigger the stimulus (e.g. right / left hand). Here we examined this issue in the visual domain using behavioral and neural measures (fMRI). Healthy participants judged the relative brightness of identical visual stimuli that were either self-triggered (using right or left hand button presses), or triggered by the computer. By presenting stimuli to either the right or left visual field, we biased visual-evoked responses to left / right visual cortex. We found stronger perceptual modulations when the triggering hand was ipsi (rather than contra) lateral to the stimulated visual field. At the neural level, we found that despite identical physical properties of the visual consequence, evoked fMRI responses in right and left visual cortices differentiate the identity of the triggering hand (left / right). Our findings support a model in which voluntary actions induce sensory modulations that follow the anatomical-functional bias of the motor system.

## Introduction

Perception is a process that does not depend solely on the physical properties of the stimulus but rather on complex interactions between those physical properties and the neural state of the perceiver. Therefore, the same stimulus can be perceived differently each time, depending on context. For example, when presented with bi-stable stimuli (such as the Rubin vase-face illusion or the Necker cube), perception fluctuates over time although the physical properties of the stimulus remain unchanged (Hesselmann et al., 2008; Iemi et al., 2017). Modulations of neural states, and subsequent perception, have been shown to depend on various contextual variables such as attention (as in the cocktail party effect; Arons, 1992), stimulation history (first vs. repeated stimulation; Grill-Spector et al., 2006; Krekelberg et al., 2006), and expectancy (Näätänen and Kreegipuu, 2011; Todorovic et al., 2011).

An important factor that has been shown to shape the neural state in sensory regions, and perception of sensory stimuli, is voluntary movement (Schütz-Bosbach and Prinz, 2007; Hughes et al., 2013; Reznik and Mukamel, 2018). Previous studies have shown that when sensory stimuli are the consequence of voluntary movement, evoked neural responses and perceptual reports are modulated relative to neural and perceptual responses to identical stimuli triggered by an external source (Hughes et al., 2013). A classic example for this effect comes from the tactile domain, where self-initiated (vs. externally initiated) tactile stimuli are perceived as less ticklish (Blakemore et al., 1999), and evoke less activity in primary somatosensory cortex (Blakemore et al., 1998). Similar modulations were found in the auditory (Baess et al., 2009; Lange, 2011; Reznik et al., 2015a) and visual domains (Stenner et al., 2014; Yon and Press, 2017; Mifsud et al., 2018). Notably, the direction of modulation by voluntary movement is inconsistent, with some studies reporting attenuation of responses, (Blakemore et al., 1998; Weiss et al., 2011; Dewey and Carr, 2013), and others reporting enhancement (Hughes and Waszak, 2011; Ackerley et al., 2012; Reznik et al., 2014). In the visual domain, behavioral and neural modulations have been reported with respect to perceived stimulus intensity (Cardoso-Leite et al., 2010; Mifsud et al., 2016; Yon and Press, 2017; Csifcsak et al., 2019), movement speed and direction (Dewey and Carr, 2013; Desantis et al., 2014), and detection of temporal delays (Matsuzawa et al., 2005; Benazet et al., 2016; van Kemenade et al., 2016).

With respect to the underlying mechanism, it has been suggested that modulations of self-triggered sensory stimuli are driven by copies of the motor commands that are sent from the motor system during voluntary movement (“efference copies”) to sensory regions that are expected to process their upcoming sensory consequences (Wolpert et al., 1995; Wolpert and Miall, 1996). Such efference signals are believed to modulate the neural state in the relevant sensory regions, resulting in differential processing of the actual reafferent (sensory) signal when it finally arrives. Efference copies have been suggested to play an important functional role in various domains including sense of agency (Gentsch and Schutz-Bosbach, 2011; Burin et al., 2017; Haggard, 2017). However, despite important basic and clinical implications ascribed to such efference signals (Pynn and DeSouza, 2013; Shergill et al., 2014), their underlying source and mechanism is poorly understood.

A prominent feature of the motor system is its hemispheric laterality bias in which control of an effector on one side of the body is associated with neural activity predominantly biased to one hemisphere. At the cortical level, it is mostly the contralateral hemisphere, while in the cerebellum, it is mostly associated with the ipsilateral cerebellar hemisphere (Kalaska and Rizzolatti, 2013). Given the premise that the source of efference copies resides within the motor system generating the action, it is plausible that a similar hemispheric bias will be observed in the magnitude of sensory modulations. Indeed in the auditory modality we have recently reported stronger perceptual / neural modulations when active motor and auditory cortices reside within the same (rather than across) hemispheres (Reznik et al., 2014). A hemispheric bias in sensory modulations at the behavioral and neural levels, that is compatible with the known anatomical-functional bias of the motor system, would provide important insight with respect to the underlying mechanism of such signals. In the current study, we examined the hemispheric bias of sensory modulations in the visual domain, using behavioral and neural measures (fMRI) in healthy participants. To this end, we manipulated the relationship between the stimulated visual field (right vs. left visual field), causal agent generating the stimulus (self / external), and identity of the effector participants used to trigger the stimulus (right / left hand).

## Methods

### Participants

Thirty-three participants, naïve to the purposes of the study, were recruited. All participants were healthy, right handed and had normal or corrected to normal vision. At the first behavioral session, data from five participants were excluded from further analysis due to low performance on the behavioral task, leaving data from twenty-eight participants (11 males, mean age 24.41, range 18-30 years).

Only participants who successfully completed the behavioral part of the study (see below) were asked to continue to an fMRI session. Two participants out of 28 requested to terminate the fMRI experiment before data collection was completed and four other participants had large head movements during the scan and therefore were excluded from the fMRI analysis, leaving fMRI data from twenty-two participants (9 males, mean age 24.23, range 18-30 years).

The study conformed to the guidelines that were approved by the ethical committee in Tel-Aviv University and the Helsinki committee of the Sheba medical center. All participants provided written informed consent to participate in the study and were compensated for their time.

### Behavioral session

In order to assess sensory modulation of self-generated visual stimuli, participants were engaged in a two alternative force choice (2AFC) task regarding the brightness level of two identical visual stimuli triggered either by the participant or the computer.

In each trial, participants were presented with two visual stimuli consecutively (passive / active). A trial began with a change in the color of the fixation point from dark gray to white, which cued the appearance of the first (passive) visual stimulus 500ms later. The stimulus was a gray circle, 2.5° in diameter that appeared for 400ms either 2.5° to the right or to the left of a fixation point (1°X1°) (see Figure 1A). After the passive stimulus disappeared, participants pressed a button in order to trigger the second (active) stimulus. Participants were instructed which hand to use (right / left) at the beginning of each block. After the active stimulus disappeared, participants were requested to report which stimulus was brighter, by pressing one of two buttons with the opposite hand to the one they used to trigger the active stimulus. Participants were instructed to answer as best as they can, and to guess if they can’t see any difference between the stimuli. Unbeknownst to the subjects, in 90% of trials both stimuli (passive / active) were identical and in the remaining 10% “catch trials” there was a real difference of 30% in brightness. These “catch trials” were inserted in order to ensure participants were attending the stimuli and performing the task. Inter-trial interval was randomized between 1-3 seconds. All stimuli were presented on a 24’ screen using Psychtoolbox-3 (www.psychtoolbox.org) on MATLAB 2015b (The MathWorks, Inc., Natick, Massachusetts, United States). This procedure was similar to the one applied by Reznik et. al. (2015b) in the auditory modality.

**Figure 1:**
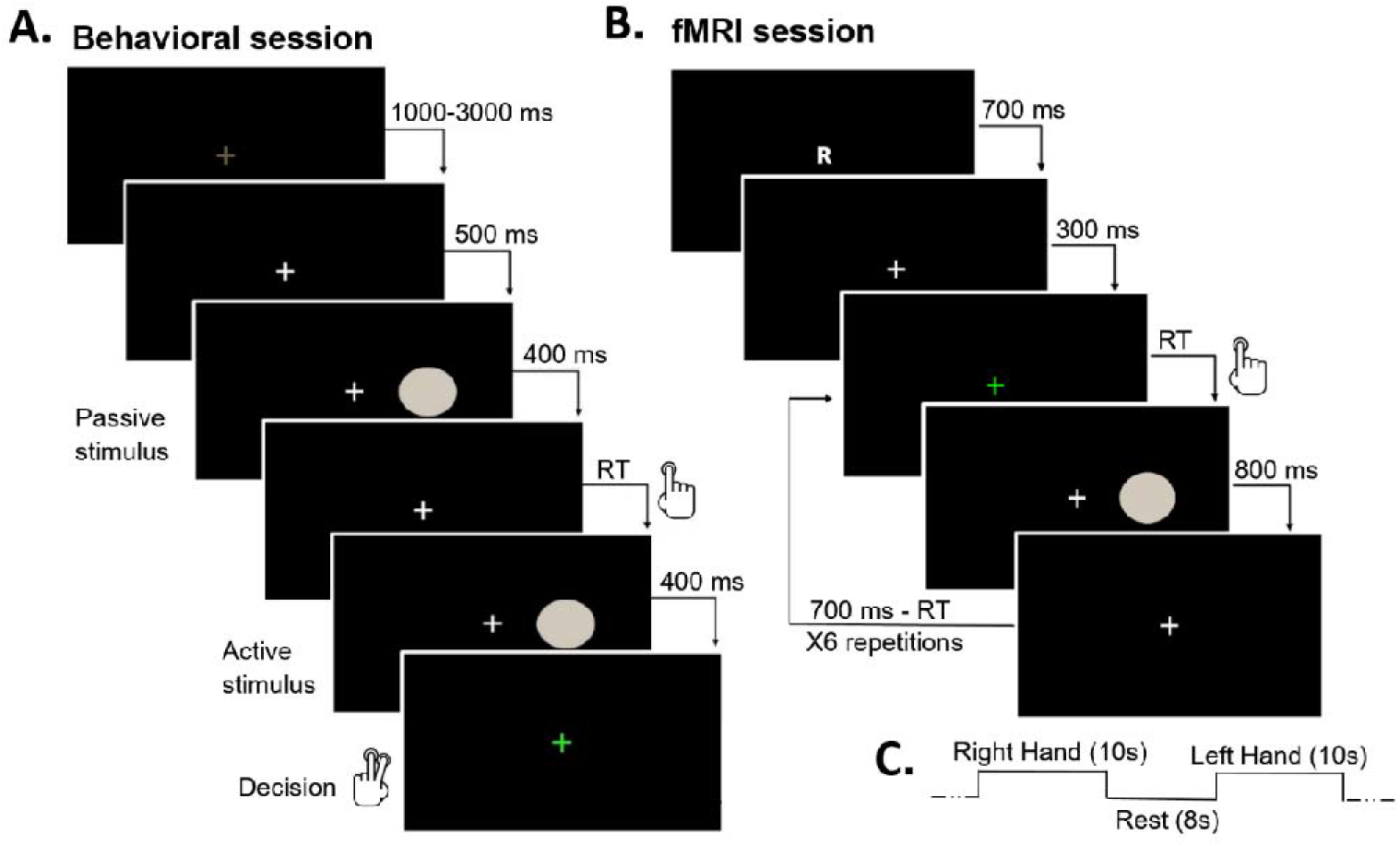
experiment procedure for behavioral and fMRI sessions. Figure 1-A. Behavioral session design: example of a single trial from a right-hand right visual-field block. Participants reported which stimulus was brighter using the other hand (left in this example). B. fMRI session design: Example from a right-hand block in a right visual-field run. C. Block design scheme for fMRI experiment. Within each run, the order of Right and Left hand blocks was randomized while stimulated visual field was kept constant.

Participants performed a total of 4 blocks corresponding to the two visual fields and two hands used to trigger the active stimulus (one block per condition). Each block contained 60 trials (6 of which were “catch trials”). Before each block, participants were informed about the stimulated visual field and which stimulus-triggering hand to use. These were kept constant throughout the block. Order of blocks was counter-balanced across participants. Participants with accuracy rates lower than 75% in the “catch trials” were disqualified from further analysis on the grounds of poor performance. Participants were informed they will receive feedback on their performance relative to previous participants after two blocks and at the end of the experiment (“Above Average” or “Below Average” performance). Feedback was based on the “catch trials” during the experiment. “Above Average” feedback was given to all participants with accuracy rates above 75%.

Before each block, participants went through 12 trials of training in order to establish the mapping between button press (right / left hand) and sensory outcome (right / left visual field). In these training trials, participants had a total of 4 “catch trials”, in which they received negative feedback if they answered incorrectly (by a red ‘X’ appearing on the screen). Participants did not receive error feedback on the identical trials during training. This was done in order to maintain each participant’s inherent selection bias. No feedback was provided during experimental trials.

Eye-tracking data were recorded using SMI RED-m 500Hz eye-tracker (SensoMotoric Instruments GmbH, Germany), and was analyzed in real-time using iViewX API for MATLAB. In the eye-tracker calibration procedure, eye-tracking accuracy was kept below 0.7°. Participants were instructed to fixate on the fixation cross throughout the experiment, and were informed that breaking fixation will result in trial disqualification and the addition of another trial to the block. Any eye movement larger than 1.5° from the fixation point during stimulus presentation triggered a “fixation break” screen, followed by the re-initiation of the trial. Due to technical problems, eye-tracking data from two participants was not obtained.

### fMRI session

fMRI session included 8 functional runs and one anatomical run. Throughout all functional runs participants were requested to fixate on a cross (1°X1°) in the middle of the screen. The first two functional runs were used to localize visual areas associated with right and left visual fields. During these runs, participants were instructed to fixate, while a flickering checkerboard appeared either on their right or left visual field. The checkerboard flickered at 10Hz and covered half the monitor (size 12.6°X 11.2°). The checkerboard was presented for 6s and then disappeared for 8s of rest, repeating for a total of 16 times in each run (8 per visual field). The order of right and left visual field stimulation was randomized. We used two identical runs instead of one long run in order to minimize fixation breaks due to participants’ fatigue.

The other six functional runs were designed to examine differential activity evoked by visual stimuli triggered with the right vs. left hand (experimental runs). These runs were organized in a block design, consisting of 10s blocks separated by 8s of resting period, during which a fixation cross appeared on a black screen. Each run consisted of 16 blocks, 8 per triggering hand. In each experimental run the stimulated visual field was kept constant (either right or left visual field condition), while the triggering hand changed across blocks. Overall, participants performed three runs for each visual field. Participants were informed that during the run they will be instructed to press a button either with their right or left hand and that their presses will trigger a visual stimulus, a gray circle similar to the ones presented on the behavioral session, either on the right or left visual field. Similar to the behavioral paradigm, visual stimuli were 2.5° in diameter and appeared 2.5° either to the right or left of a fixation cross. Stimuli were presented on a 32’ monitor and viewed by the participants through a mirror placed on the MRI head coil.

Each block started with a 700ms presentation of either the letter ‘R’ or ‘L’ (0.5°X0.5°) on the center of the screen, indicating the hand to be used for triggering the visual stimulus during the block (right or left respectively). After the letter disappeared, participants were instructed to press with the appropriate hand as fast as possible every time the fixation point changed color to green (once every 1.5s; see figure 1B). Prior to each experimental run, participants were informed that each button press will trigger a visual stimulus in a specific location (which was kept constant throughout the run: either right or left visual field). Overall, participants triggered 6 visual stimuli on each block. Each stimulus was presented for 800ms immediately following button press. If the participant’s RT was longer than 700ms, a red ‘X’ (1.4°) appeared on the screen, indicating slow responses, and the entire block was removed from further analysis. This measure was taken in order to maintain a constant presentation pace in both Right and Left hand conditions and to make sure blocks were precisely timed to TR. Order of right and left hand blocks within a run was randomized.

In order to keep participants attentive to the visual stimuli, in 2-4 of the blocks one of the circles was blue instead of gray. Participants were requested to count how many times they saw a blue circle throughout the run and verbally report it at the end of each run. Blocks with blue circles were removed from further analysis.

Throughout the experiment, participants’ eye movements were monitored in order to ensure fixation. Eye-tracking data were collected using an MR-compatible Eyelink ® 1000 plus (SR Research Ltd., Mississauga, Ontario, Canada), sampled at 500Hz. Eye-tracking calibration accuracy was kept below 1°.

### fMRI data acquisition

Functional imaging was performed on a Siemens Magnetom Prisma 3T Scanner (Siemens Healthcare, Erlangen, Germany) with a 64-channel head coil at the Tel-Aviv University Strauss Center for Computational Neuroimaging. In all functional scans, an interleaved multi-band gradient-echo echo-planar pulse sequence was used. 66 slices were acquired for each volume, providing whole-brain coverage (slice thickness 2mm; voxel size 2^3^mm; TR= 2000ms; TE=30ms; flip angle=82°; field of view=192mm; acceleration factor=2). For anatomical reference, a whole-brain high-resolution T1-weighted scan (slice thickness 1mm; voxel size, 1^3^mm; TR=2530ms; TE=2.99ms; flip angle=7°; field of view=224mm) was acquired for each subject.

### Behavioral data analysis

In order to evaluate the behavioral magnitude of sensory modulation, a modulation index was calculated for each combination of triggering hand and stimulated visual field. This index was defined as the absolute difference between the proportion of trials in which the subject chose the self-triggered stimulus as brighter, and chance level of choosing either stimulus as brighter (active / passive; 0.5), as in the formula below:

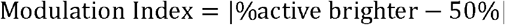

This measure represents the deviation from chance for each participant to report the stimulus from one condition (active / passive) as brighter. A modulation index of 0 indicates an equal proportion of trials in which the active or passive stimulus was reported as brighter. Note, this index is non-directional, and emphasizes the magnitude of deviation irrespective of the tendency to report, for example, the active condition as brighter (or vice versa). Importantly, we used this index to compare changes in report tendency across conditions rather than examine general tendency biases to report one condition as brighter. This index was used as the dependent variable in the analysis of the behavioral session. Behavioral data were analyzed using a 2X2 repeated measures ANOVA with stimulated visual field (right vs. left) and stimulus triggering hand (ipsilateral vs. contralateral to stimulated visual field) as independent variables. Analysis was performed using JASP (JASP Team, 2019. Version 0.10.1).

### fMRI data analysis

fMRI data preprocessing and first level GLM analysis was conducted using The FMRIB’s Software Library’s (FSL v5.0.9) fMRI Expert Analysis Tool (FEAT v6.00) (Smith et al., 2004b). The data from each experimental run was brain-extracted, slice-time corrected, high-pass filtered at 100s (0.01Hz), motion-corrected to the middle time-point of each run, smoothed with a 5 mm FWHM kernel, and corrected for autocorrelation using pre-whitening (as implemented in FSL). We excluded from further analysis participants with more than one run during which the absolute displacement values exceeded 2 mm. All images were registered to the high-resolution anatomical data using boundary-based reconstruction and normalized to the Montreal Neurological Institute (MNI) template using nonlinear registration.

Localizer data were analyzed using a general linear model with two regressors - right visual field and left visual field. A conventional double gamma response function was convolved with each of the regressors in order to account for the known lag of the hemodynamic responses. Additionally, the six motion parameter estimates from the rigid body motion correction were included in the model as nuisance regressors. We calculated both right visual field>left visual field and left visual field>right visual field contrasts. Results from these contrasts were Bonferroni corrected for multiple comparisons with α=0.05.

The six experimental runs were analyzed using a multi-voxel pattern analysis (MVPA) classifier approach. We used a Java implementation of a support vector machine (SVM) classifier (Chang and Lin, 2011) to discriminate right and left hand activation patterns in the visual cortex. For each voxel and each block, we calculated percent signal change of the last TR (TR=5), relative to time course mean. This resulted in a total of 36-40 values for each voxel in each participants’ brain (18-20 for Right-Hand condition and 18-20 for Left-Hand condition in each visual field; The exact number of values within participant was identical in both conditions, however there were slight differences between participants due to response errors in the task). For each voxel, defined as center-voxel, we outlined a neighborhood which included the center voxel and its 26 closest voxels (in Euclidean distance).

To estimate classification level between right and left hand blocks within a certain visual field, we used an SVM classifier with a linear kernel, and a leave-one-block-out approach. For each neighborhood we used 250 iterations of leave one block out and used the averaged accuracy level across all iterations as the decoding accuracy of the center voxel. In order to determine the significance level of our classification values, we used permutation analysis to create a shuffle distribution for each neighborhood of voxels. For each participant, we shuffled the data labels (right / left hand blocks) and repeated the same analysis that was performed on the real data. Overall, for each participant we calculated a map of real data accuracy-levels and 100 maps of accuracy-levels based on shuffled data. To determine group-level significance, we used the permutation scheme suggested by Stelzer et al. (2013). First, we averaged all the real accuracy-level maps across subjects to create a group average map. Next, we randomly chose one shuffle map from each participant and averaged those shuffled maps across participants to create one average shuffled map. This procedure was repeated 10000 times for the shuffled maps, providing a distribution of shuffled data accuracy maps. Thus, the minimal p-value of the real map is 0.0001. The p-values obtained from this procedure were corrected for multiple comparisons using false discovery rate approach (Benjamini and Hochberg, 1995) with q=0.05.

### Functional connectivity analysis

In order to examine functional connectivity between visual and motor cortices, and between visual cortex and the cerebellum, we calculated the correlation between the time courses in those regions. For each participant, we defined separately the cerebellum, motor and visual regions in both hemispheres. Visual regions of each participant were defined by GLM contrast of each visual field condition’s experimental runs, using the contrast (Right hand + Left hand) > rest, in order to ensure we used areas responsive to the visual stimulus regardless of triggering hand. In order to restrict our ROI to visual cortex, the results of this contrast were intersected with the visual localizer. For correlation analysis we used the averaged time course from the most significant voxel in the ROI and it’s 26 nearest neighbors. Motor regions and cerebellum of each participant were defined using a GLM contrast of each visual field condition’s experimental runs, using the contrasts Right hand > Left hand and Left hand > Right hand. We intersected the results from these contrasts with the anatomical masks of right and left motor cortex taken from the Harvard-Oxford lateralized cortical structural atlas (Desikan et al., 2006), and with a cerebellum mask taken from the MNI structural atlas (Mazziotta et al., 2001). As in the visual cortex, for correlation analysis we used the averaged time course from the most significant voxel in the ROI and it’s 26 nearest neighbors. Next, for each visual cortex time course we calculated the Pearson correlation with the time course of left and right motor cortex separately, and left and right cerebellum separately. To compare between correlations of each stimulated visual cortex with ipsilateral and contralateral motor cortex and cerebellum across subjects, we used a paired t-test applied on the Fisher’s transformation of the correlation values.

## Results

### Perceptual modulation - behavioral results

In order to examine the influence of stimulus-triggering hand on sensory modulations, we used a 2×2 repeated measures ANOVA (*N*=28), with visual field and triggering hand (ipsilateral or contralateral to visual field) as independent variables. We found a significant main effect for triggering hand (*F*(1,27)=4.85, *p*=0.03; see figure 2), indicating higher modulation index for stimuli triggered with the hand ipsilateral to the stimulated visual field (*M*=0.16, *SD*=0.07), relative to stimuli triggered with the hand contralateral to the stimulated visual field (*M*=0.13, *SD*=0.08). We did not find a significant main effect of stimulated visual field (right visual field: *M*=0.14 *SD*=0.10; left visual field: *M*=0.15 *SD*=0.09; *F*(1,27)=0.80, *p*=0.38), or a significant interaction between stimulated visual field and triggering hand (*F*(1,27)=0.153, *p*=0.699).

**Figure 2:**
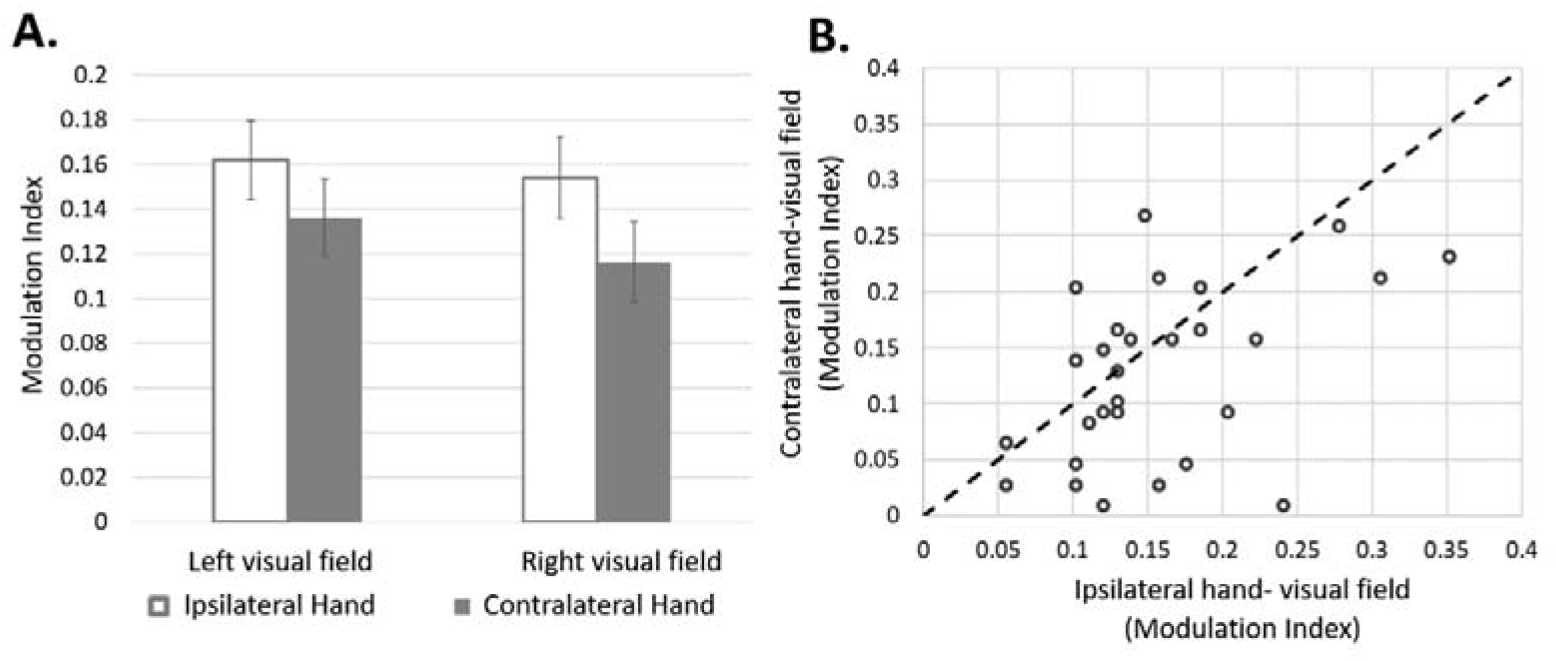
Hand dependednt modulation of perception. Figure 2-A. Perceptual modulations. Modulation index (group results, *N*=28) according to visual field of action consequences. A significant main effect for laterality of triggering hand (ipsilateral vs contralateral to the stimulated visual field) (*F*(1,27)=4.85, *p*<0.05). Error bars represent SEM. B- Modulation index of individual subjects according to laterality of triggering hand (ipsilateral/contralateral), collapsed across visual fields. Each dot represents the averaged modulation index of each participant and dashed line represents equal modulation for ipsilateral/contralateral conditions.

Comparing reaction times (RTs) for triggering the active stimulus, we did not find a main effect for visual field (Right visual field: *M*=0.67s, SD=0.40; Left visual field: M=0.64s, *SD*=0.29; *F*(1,27)=0.38, *p*=0.55), and no main effect for triggering hand (ipsilateral hand: *M*=0.64s, SD=0.27; contralateral hand: M=0.66s, *SD*=0.31; *F*(1,27)=0.35, *p*=0.56). Also, we did not find a significant interaction effect between visual field and triggering hand (Right visual field: ipsilateral hand - *M*=0.62s, *SD*=0.28, contralateral hand - *M*=0.73s, *SD*=0.52; Left visual field: ipsilateral hand - *M*=0.67s, *SD*=0.32, contralateral hand - *M*=0.60s, *SD*=0.26; *F*(1,27)=3.10, *p*=0.09). Average performance on “catch trials” across participants was at ceiling level in all conditions (mean accuracy = 95.24% across all conditions).

### Neural modulations - fMRI results

#### Visual ROIs

In order to functionally define the visual cortex, we first performed a visual localizer task (see methods). We used GLM with a contrast of Right Visual Field>Left Visual Field to define regions sensitive to visual stimulation in either right or left visual fields. Figure 3A shows a multi-subject (*N*=22), MNI normalized, Boolean map of voxels showing significant voxels, either positive (red) or negative (blue), for this contrast (p<0.05 Bonferroni corrected; 3,852 voxels). This map was used as a mask to define visual regions in the experimental runs.

**Figure 3:**
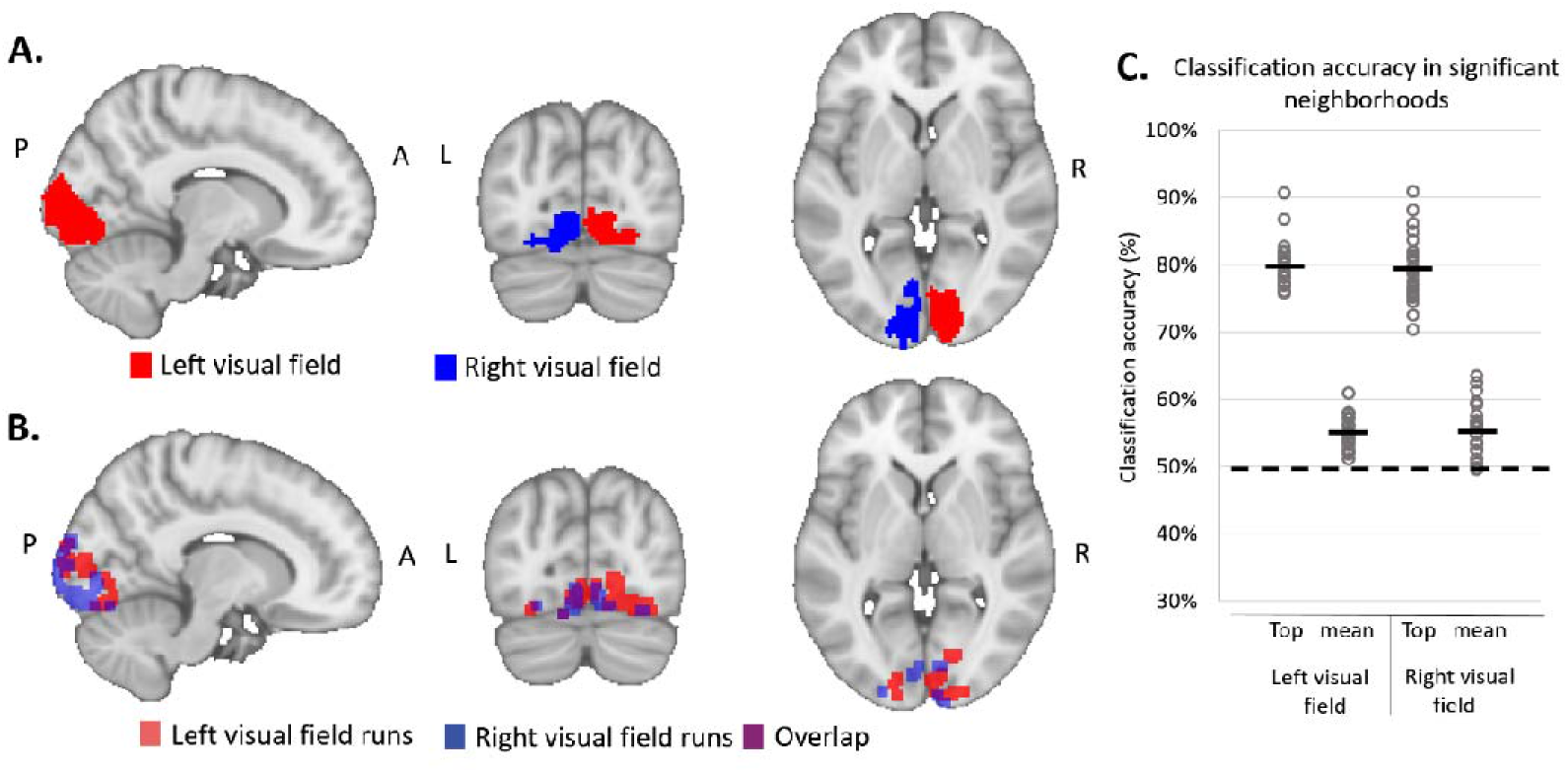
hand dependednt modulations of fMRI signals. Figure 3-A. Boolean map showing significant voxels in the visual localizer runs (GLM analysis, *n*=22; <0.05 Bonferroni corrected; see methods section). Red voxels correspond with significant voxels in the contrast left visual field>right visual field and blue voxels for the opposite contrast. B. Classification analysis (group results). Boolean map of significant voxel neighborhoods separating right and left hand in visual cortex, despite identical visual stimuli (*p*<0.05 FDR corrected; see methods). Red voxels correspond with voxels from left visual field runs and blue voxels correspond with voxels from right visual field runs. Purple areas represent overlap between voxel neighborhoods across runs. Note the significant discrimination of hands in both hemispheres irrespective of stimulated visual field. P – posterior, A – anterior, R – right hemisphere, L- left hemisphere. C. Individual subjects’ classification accuracy levels in significant voxels for each stimulated visual field condition. Significant voxels were identified from the group analysis (see methods). Top represent each subject’s voxel with the highest accuracy in the significant voxels in visual cortex, while mean represent each subject’s mean classification accuracy across all significant voxels. Dashed line represents chance accuracy level (50%).

### Neural modulations in visual cortex according to triggering hand - SVM results

In order to examine differential modulation of visual cortex, we classified fMRI activity patterns evoked by identical visual stimuli according to triggering hand (right / left). Our group analysis revealed neighborhoods of voxels significantly distinguishing between the two hand conditions in each visual field (right / left) (*p*<0.05 FDR corrected; Figure 3B). This differentiation was found in both visual cortices, regardless of the stimulated visual field (see figure 3B). In the Right visual field condition, we found 206 significant neighborhoods, 60 of which were in the left (contralateral) visual cortex. In the Left visual field condition, we found 268 significant neighborhoods, 149 of which were in the right (contralateral) visual cortex. 23 neighborhoods overlapped between right and left visual field conditions (for mean and individual decoding accuracy levels across subjects see figure 3C).

### Connectivity between stimulated visual cortex and motor regions

In order to examine whether motor regions exert stronger modulations on sensory regions residing within the same hemisphere, we calculated the correlations between activity in each visual cortex with activity in right / left motor cortices and right / left cerebellum. We found that in left visual field runs, activity in the right visual cortex was more strongly correlated with the right (within hemisphere) motor cortex (*r*=0.32) than with the left (across hemisphere) motor cortex (*r*=0.27; *t*=2.29, *p*=0.03). In Right visual field conditions, we found no significant difference in correlation between Left visual cortex and left (within hemisphere; *r*=0.23) or right (across hemisphere; *r*= 0.25) motor cortices (*t*=0.81, *p*=0.43; See figure 4). a similar connectivity analysis performed between stimulated visual cortex and right / left cerebellum showed no significant effect. In left visual field condition, connectivity between right visual cortex and right (*r*=0.18) or left (*r*=0.21) cerebellum (*t*=1.46, *p*=0.16); In right visual field condition, connectivity between left visual cortex and left (*r*=0.18) or right (*r*=0.18) cerebellum (*t*=0.11, *p*=0.91).

**Figure 4:**
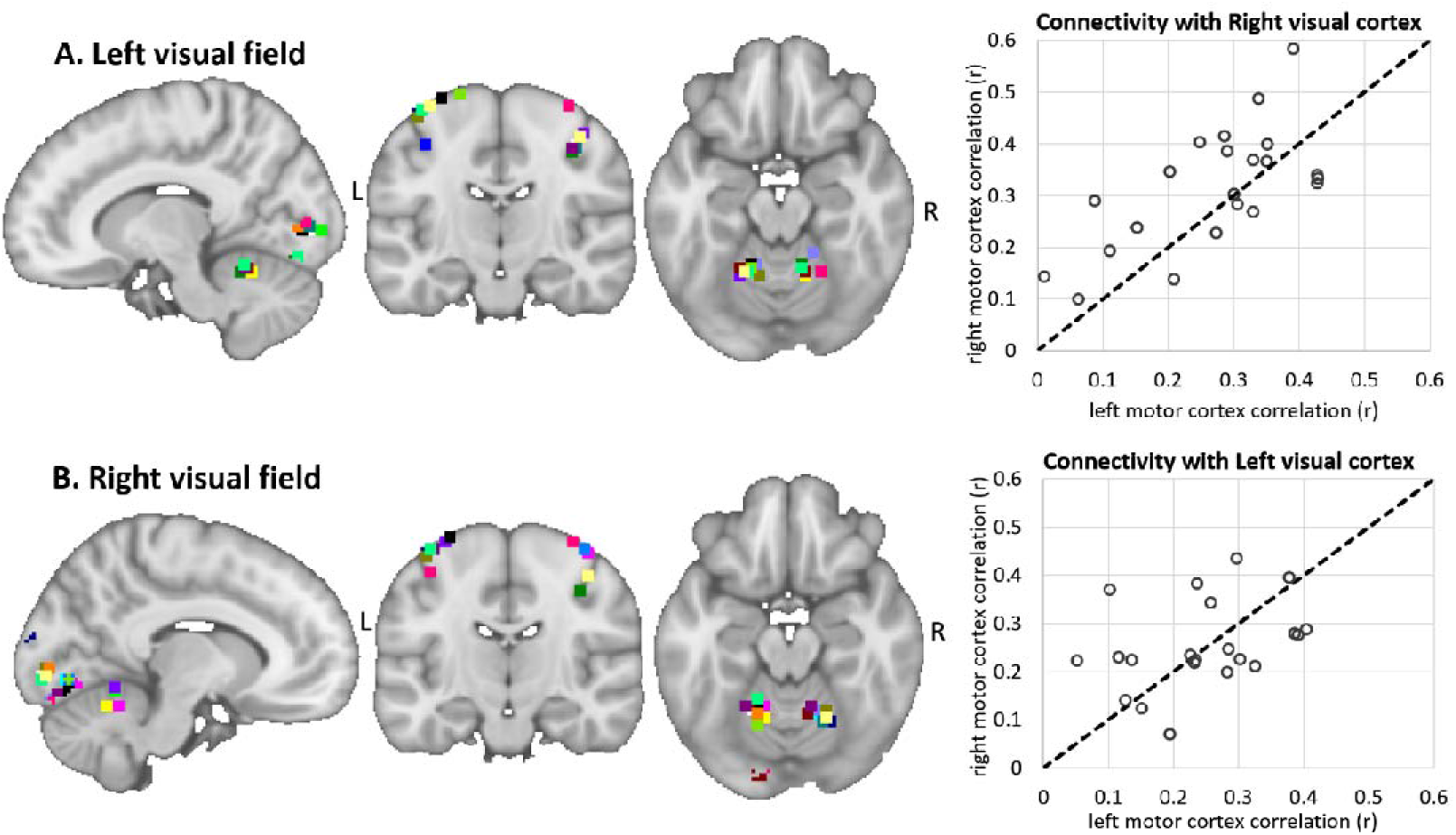
functional connectivity between visual cortex and motr regions. Figure 4- functional connectivity between stimulated visual cortex and right/left motor cortices during task execution. Visual, motor, and cerebellar ROIs of individual subjects (*N*=22) presented on sagittal, coronal and axial views of an MNI template (left panels; each color represents a 27 voxel ROI of an individual subject; see methods). Panel A corresponds with left visual field runs and Panel B with right visual field runs. Scatter plots to the right display the functional connectivity (Pearson’s r) between right (top) or left (bottom) visual cortex and the two motor cortices. Each dot represents data of a single subject, dashed line indicates equal functional connectivity with right/left motor cortices. During left visual field runs (Panel A), right visual cortex exhibited stronger functional connectivity with right vs. left motor cortex (t=2.29, p<0.05) but during right visual field runs (Panel B) no significant difference was found (t=0.81, p=0.43). R – right hemisphere, L- left hemisphere.

## Discussion

In the current study, we examined differences in perceptual and physiological responses to identical visual stimuli while manipulating the stimulus-triggering hand. In agreement with our hypothesis, our behavioral findings show stronger perceptual modulation for stimuli triggered with the hand ipsilateral (vs. contralateral) to the stimulated visual field. Our fMRI results show that despite identical physical properties of the visual stimulus, neural activity in visual cortex differentiated the stimulus-triggering hand. Finally, functional connectivity analysis between visual cortex and ipsilateral / contralateral motor cortex and cerebellum, showed stronger connectivity between right visual cortex and the ipsilateral (right) motor cortex, and no difference in connectivity strength with right / left cerebellum. For left visual cortex, no differences in functional connectivity were found.

Results from our SVM analysis show that visual cortex responds differently to identical visual stimuli, depending on the stimulus-triggering hand. In other words, actions that generate visual feedback, are associated with responses in visual cortex that are not only sensitive to the visual properties of the stimulus, but also to the hand that was used to generate it. This non-intuitive result is supported by recent EEG evidence showing differential magnitudes of sensory attenuation (expressed in the evoked response signal) for stimuli generated with various effectors (e.g. the hand, eyes, or mouth) in the visual (Mifsud et al., 2018) and auditory (Mifsud et al., 2016) modalities. In addition, previous fMRI studies report networks in which preparatory neural activity preceding movement onset differentiates the limb to be used (Gallivan et al., 2013; Gallivan et al., 2015). Importantly, these fMRI studies report that during the planning phase, neural activity in sensory regions shows limb specificity (Gallivan et al., 2019). Thus, together with our results, there is sufficient evidence demonstrating that signals in visual cortex carry limb-specific information about motor commands during action planning and execution.

Interestingly, despite presenting visual stimuli in one visual field in a given experimental condition, we found that limb-dependent modulations were not restricted to the visual cortex contralateral to the stimulated visual field. Instead, we find significant modulations also in ipsilateral visual cortices. One interpretation of this finding is that button presses modulate signals in visual cortex in a non-specific manner, including visual regions that are not engaged by the stimulus. An alternative explanation is that button presses modulate evoked responses in ipsilateral visual cortex. Although it is common to associate visual stimulation in one visual field with activations in the contralateral primary visual cortex (Gilbert, 2013), previous studies have shown activations also in the visual cortex ipsilateral to the stimulated visual field (Tootell et al., 1998; Smith et al., 2004a). Therefore, in light of these findings, the hand-dependent modulations we report in both visual cortices may still reflect specific motor signals to sensory regions that are sensitive to the visual stimulus. In the current experiment, we found significant ipsilateral activations during the localizer runs supporting specificity of modulations. However, in the experimental runs, we failed to see significant ipsilateral activations at the group level, perhaps due to lower salience of the visual stimulus used (gray circle vs. checkerboard) or poor alignment of weak ipsilateral activations across subjects. Therefore, based on the results of the current study, we cannot resolve the degree of specificity of the motor modulations.

Related to this point, it is not clear at this stage whether the modulations we report in visual cortex are due to the causal link between the button presses and the visual consequences. Since the aim of the current study was to explore the dependency of sensory modulations on limb identity, we did not manipulate the causal link, for example, by measuring neural responses in visual cortex during right / left button presses with no visual consequences. To the best of our knowledge, no previous study reported limb-specific modulations of visual cortex in the absence of visual stimulation. The forward model framework would suggest that expectancy of sensory consequences plays an important role in sensory modulations. Therefor under this framework, in the lack of expected visual consequences, differential hand-dependent modulations in visual cortex should not be found. Whether limb-specific modulations in visual cortex can be found in the absence of visual stimulation is an interesting question for a future study.

While it is assumed that the efference copies originate from the motor system, as proposed by the forward model (Wolpert et al., 1995; Crapse and Sommer, 2008), the anatomical origin of such signals is not known. Some studies suggest that efference copies originate from the active motor cortex (Gandolla et al., 2014; Reznik et al., 2014), while others suggest the cerebellum to be the origin of these signals (Blakemore et al., 2001; Person, 2019). By manipulating the hands used to generate the stimulus, we engaged different motor pathways. Similarly, by manipulating the stimulated visual field we engaged different visual cortices. Therefor to the extent that active motor pathways are the source of sensory modulations, it is plausible for modulations in sensory cortex to reflect their different motor origins. Our functional connectivity data demonstrate that right visual cortex is more strongly connected with the right (vs. left) motor cortex, while functional connectivity with right / left cerebellum was not significantly different. While these results are in better agreement with a cortical source of modulations, it is important to note that the hemodynamic response function as measured by the MR scanner is different between the cerebral cortex and the cerebellum (Hossein-Zadeh et al., 2003; Chen and Desmond, 2005), a fact that may potentially bias our functional connectivity analysis in favor of cortical regions (visual / motor) over cortex-cerebellum connectivity. In the left visual cortex, we found no significant differences in functional connectivity with either right / left motor cortex or right / left cerebellum suggesting possible hemispheric differences in connectivity. These issues remain to be resolved in future studies.

There is an ongoing discussion regarding the functional role of efference copies and sensory modulations, including agency attribution, and desensitization of sensory apparatus (Gentsch and Schutz-Bosbach, 2011; Burin et al., 2017; Haggard, 2017). Although our study does not address these functional roles, our results suggest that to the extent that sensory modulations are involved in such processes, these processes should have a component of limb specificity. Additionally, it should be noted that to date, there is no direct causal evidence linking the behavioral and physiological phenomena of sensory modulations, and many studies report either one. Although in the current study we report both measures from the same subjects, differences in design and analysis levels precluded us from performing a direct comparison of measures across subjects.

A large body of literature reporting sensory modulations in humans has focused on the auditory and tactile modalities (Blakemore et al., 1999; Baess et al., 2009; Lange, 2011; Weiss et al., 2011; Reznik et al., 2015b), with fewer studies characterizing this phenomenon in the visual domain (Cardoso-Leite et al., 2010; Straube et al., 2017; Yon and Press, 2017; Csifcsak et al., 2019). Thus our behavioral results provide an expansion of the current literature in the visual domain with respect to stimulus brightness. Moreover, to the best of our knowledge, there is no previous evidence demonstrating effector-dependent sensory modulations at the behavioral level. A common finding in previous behavioral results, irrespective of sensory modality, is attenuation of reported perceptual intensity (e.g. tactile pressure or sound amplitude; Blakemore et al., 1999; Reznik et al., 2015b). In our current results, the direction of perceptual modulations was not consistent across subjects, with some subjects reporting increased stimulus brightness and others reporting decreased stimulus brightness of the self-generated stimuli. Nevertheless, when comparing the magnitude of modulations across hands, we found stronger modulations when the stimulus-triggering hand and stimulated visual field are ipsilateral and thus predominantly processed in the same hemisphere. We recently proposed a model in which sensory regions are more strongly modulated when the motor region engaged in producing the action resides in the same hemisphere (Reznik et al., 2014). The behavioral and neural results of the current study are in agreement with such a model.

In motor cortex, recent evidence from EEG demonstrate differences in readiness potential depending on the coupling with a sensory consequence (Reznik et al., 2018). Thus, together with the current results, it seems that information about expected sensory consequences is stored in the motor cortex while limb-specific motor information of signal source is stored in the sensory cortex (visual cortex in the current study). Although speculative at this point, such modulations both in motor and sensory cortices may constitute an important neural loop for sensorimotor learning. In summary, by demonstrating limb-specific sensory modulations at both the behavioral and neural levels, our results help constrain future models describing their underlying mechanisms and provide further evidence that neural responses in regions primarily described by their sensory properties (in our case ‘visual cortex’) go beyond a simple representation of the physical / optical properties of the external world.

## Acknowledgments

This research was supported by the Israel Science Foundation (grant No. 2392/19) to R.M. The authors thank lab members for constructive comments and fruitful suggestions.

